# LAMP2A-dependent chaperone-mediated autophagy enhances oxidative stress resistance in gastric cancer cells through selective degradation of accumulated oxidized DJ-1

**DOI:** 10.1101/2025.08.24.672019

**Authors:** Shuangshuang Le, Tongtong Guo, Tianjuan Tang, Yang Zheng, Maogui Pang

## Abstract

Reactive oxygen species (ROS) function as potent activators of chaperone-mediated autophagy (CMA). While previous investigations have demonstrated that CMA facilitates cancer cell survival through selective degradation of oxidatively damaged proteins, the precise molecular mechanisms remain poorly defined. Hence, it is important to investigate the correlation between CMA and the antioxidative stress within tumor cells and elucidate its specific mechanism. LAMP2A expression profiles in gastric carcinoma cell lines and clinical specimens were quantified via qPCR, WB and immunohistochemical analysis. An in vitro oxidative stress model was established through 24-hour hydrogen peroxide (H_2_O_2_) exposure. Gastric cancer cell models with knockdown and overexpression of LAMP2A were established. Apoptotic indices and proliferative capacity were assessed through high-throughput flow cytometry and CCK-8 viability assays. The interaction between LAMP2A and DJ-1 was investigated via co-immunoprecipitation and confocal microscopy. This study demonstrates that LAMP2A is upregulated in gastric cancer and further induced by oxidative stress. Down-regulated LAMP2A impaired CMA functionality, sensitizing gastric cancer cells to oxidative cytotoxicity and significantly augmenting apoptosis rates. Elevated DJ-1 levels in gastric cancer cells are further amplified by oxidative stress, with severe stress enhancing LAMP2A-DJ-1 colocalization. Mechanistically, CMA inhibition precipitated accumulation of hyperoxidized DJ-1 isoforms, concomitant with pro-apoptotic BAX upregulation and anti-apoptotic BCL-2 downregulation. We found hyperoxidized DJ-1 as a novel CMA substrate. The LAMP2A-DJ-1 regulatory axis represents a critical adaptive mechanism whereby CMA activation maintains redox homeostasis in gastric malignancies. Specifically, CMA-mediated clearance of oxidized DJ-1 prevents pro-apoptotic protein cascade activation, thereby conferring oxidative stress resistance and promoting tumor cell survival.

## 1. Introduction

Gastric cancer (GC), ranked fifth in global cancer incidence and third in mortality [1]. While biomarkers including PD-L1, MSI, and HER2 have demonstrated therapeutic efficacy [2], their limited specificity and sensitivity underscore the urgent need for novel molecular markers to improve early diagnosis and survival outcomes. Oxidative stress is a state of imbalance between oxidation and antioxidant effects in cells [3]. Oxidative stress increases the level of reactive oxygen species (ROS) in cells. ROS function as essential regulators of physiological processes including cellular proliferation, differentiation, and apoptosis, while their supraphysiological accumulation during oxidative stress induces cytotoxic damage [4]. The excessive proliferation of tumor cells is accompanied by the production of high levels of ROS, but tumor cells can avoid aging and apoptosis caused by ROS by improving their antioxidant capacity [5]. Autophagy plays a vital role in the life activity of cells, which comprises three mechanistically distinct lysosomal degradation pathways: macroautophagy, microautophagy, and chaperone-mediated autophagy (CMA). And CMA requires lysosomal-associated membrane protein 2A (LAMP2A) to transport the substrate [6]. CMA specifically targets cytosolic proteins containing KFERQ-like motifs, which are selectively recognized by heat shock cognate 70 kDa protein (HSC70) to form a substrate-chaperone complex prior to lysosomal translocation [7]. Subsequently, the complex enters the lysosome through LAMP2A for degradation [8].

CMA activation has been implicated in tumor progression across multiple malignancies, promoting the proliferation [9]. For example, downregulation of LAMP2A can reduce the proliferation and self-renewal of glioblastoma, induce apoptosis of GSCs in vitro, and delay tumor progression in vivo [10]. LAMP2A upregulation drives colorectal cancer progression by enhancing tumor cell proliferation, invasive capacity, and apoptosis resistance [11]. Some studies have shown that inhibiting the interaction between HSC70 and LAMP2A can block CMA in NSCLC and inhibit tumor growth [12]. ROS is an activator of CMA, and CMA ensures the survival of breast cancer cells by degrading proteins produced by oxidative damage [13]. Studies have found that LAMP2A, a key receptor molecule in CMA, has an early warning effect on gastric mucosal precancerous lesions and is a specific gastric cancer marker [14]. Therefore, our study will further explore the role and molecular mechanism of CMA in regulating the anti-oxidative stress of gastric cancer cells. We hope establish a mechanistic framework for CMA-mediated oxidative stress adaptation in gastric cancer cells.

## 2. Materials and methods

### 2.1 Cell culture

Normal human gastric mucosal epithelial GES-1 cells and gastric cancer cell lines (AGS, MKN45, HGC27, MKN28) were obtained from the ATCC. Cells were cultured in DMEM (Gibco, 11995040) supplemented with 10% fetal bovine serum (FBS; Gibco, 10099-141) and 1% penicillin-streptomycin (Pen-Strep; ThermoFisher, 15140122), and maintained at 37°C with 5% CO2. All cell lines were validated through short tandem repeat (STR) profiling, with morphological and functional characteristics frequently checked, and confirmed mycoplasma-free.

### 2.2 Cell transfection

The LAMP2A coding sequence synthesized by TsingkeBiotechnology Co.,Ltd was cloned into pcDNA3.1 to generate the LAMP2A-overexpression plasmid. Cells were transfected with LAMP2A-encoding lentiviral vectors (Shanghai GeneChem Co., Ltd., PIEL248064052) at 37°C upon reaching 40% confluence.

### 2.3 RT - qPCR

Total RNA was extracted using an RNA purification kit (Invitrogen, K0731). Reverse transcription was performed with Takara reagents under standard conditions (37°C, 15 min; 85°C, 5 s). PCR amplification was conducted using a two-step cycling protocol (40 cycles): 95°C for 10 s denaturation and 60°C for 30 s extension. *LAMP2A* and *β-actin* primers (Table 1) were custom-synthesized by TsingkeBiotechnology Co., Ltd. RT-qPCR was conducted on a LightCycler480 system (Roche). *LAMP2A* and *β-actin* expression were quantified using the 2^−ΔΔ^ Ct method.

**Table 1.**
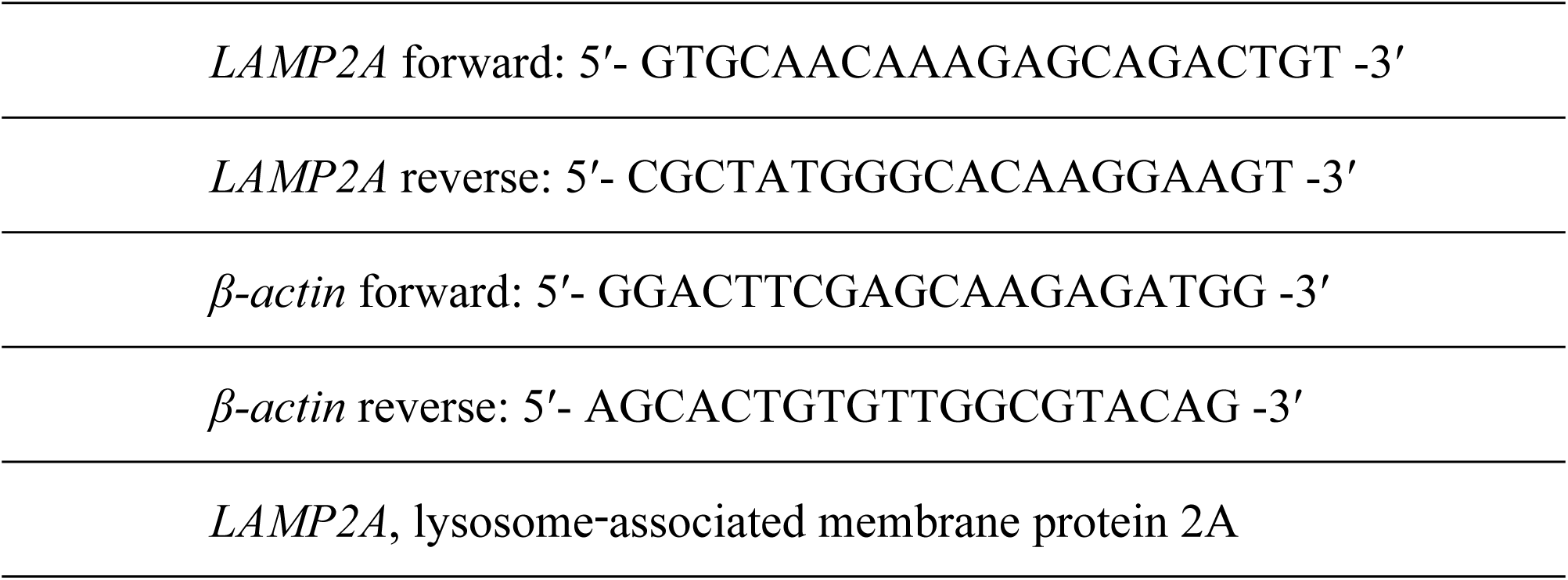
Primer and small interfering RNA sequences.

### 2.4 Cell Counting Kit-8 assay

Cells were seeded in 96-well plates (2,000 cells/well, five replicates per group) and cultured under standard conditions (37°C, 5% CO₂). Adding CCK-8 reagent (GLPBIO, GK10001) to the medium, after a 2-hour incubation period, absorbance measurements were conducted at 450 nm using a microplate reader (Thermo Fisher, VLBL00GD2).

### 2.5 Immunohistochemical staining

The gastric cancer tissues and adjacent normal tissues were purchased tissue microarrays obtained from Shanghai Outdo Biotechnology Co., Ltd (Product ID: HStmA020PG01). Tissue sections underwent sequential processing: baking at 65°C for 3 h, rehydration, EDTA-based antigen retrieval (pH 8), peroxidase blocking with 3% H_2_O_2_, and serum blocking. Primary antibody incubation used rabbit anti-LAMP2A (dilution 1:200, 28477-1-AP, Proteintech) overnight at 4°C, followed by 30 min treatment with HRP-conjugated secondary antibodies (ZSGB). DAB chromogenic reaction was performed for 1 min using a commercial kit. The tissue sections underwent hematoxylin counterstaining, followed by sequential ethanol dehydration, xylene clearing, and final mounting using neutral balsam medium. Whole-slide imaging was conducted on an Olympus VS120 system.

### 2.6 Flow cytometric analysis of apoptosis

Cells were harvested and washed twice with PBS, and resuspended in 1× Binding Buffer to achieve 1×10^6^ cells/mL. 100µL cell suspension were stained with PE Annexin V and 7-AAD (5 μL each) for 15 min in light-protected conditions.

### 2.7 Immunofluorescence

Cells were seeded in chamber slides at 5×10⁴ cells/well and cultured for 24 h. Following treatment with 1 mL 300 μM H₂O₂ for 24 h, cells were fixed with 4% paraformaldehyde (200 μL/well, 15 min RT), then washed twice with PBS (2 min/wash) and thrice with PBS (5 min/wash). Cells were permeabilized with 200 μL of 0.1% Triton X-100 (T8787, Sigma-Aldrich) for 15 min. After three PBS washes (5 min each), nonspecific binding sites were blocked with 200 μL of immunofluorescence blocking buffer (P0102, Beyotime) for 1 h at 25°C. Primary antibodies (1:200; ab199337, Abcam) were applied (200 μL/chamber) and incubated overnight at 4°C. After three PBS washes, samples were incubated with species-specific Alexa Fluor-conjugated secondary antibodies (1:500; A-11034, Thermo Fisher) for 1 h at 25°C with gentle agitation, protected from light. Nuclei were counterstained with DAPI (40728ES03, Yeasen) for 30 sec. Fluorescence imaging was performed using a Nikon A1R HD25 confocal system (AX/AXR with N SPARC). Protein localization analysis included systematic random sampling of ≥5 fields per condition, with signal specificity confirmed through negative controls lacking primary antibodies.

### 2.8 Western blotting

Following 72-hour treatment, cells were harvested and lysed in ice-cold RIPA buffer. Lysates were centrifuged at 12,000 ×g for 15 min at 4°C, and supernatants were collected. Protein concentrations were quantified using a BCA assay (23235, Thermo Fisher). Protein samples were denatured at 95°C for 5 min in a preheated thermal cycler. Electrophoresis was performed using 10% SDS-PAGE gels (4561033, Bio-Rad) at 100 V for 120 min. Resolved proteins were transferred to membranes (0.45 μm, 1620115, Bio-Rad) via wet transfer (30 V, 60 min, 4°C). Membranes were blocked with 5% non-fat milk for 1 h. Primary antibodies incubated overnight at 4°C: LAMP2A (1:1000, #Ab125068, Abcam), Dj-1(1:1000, # Ab76008, Abcam), Bcl-2 (1:1000, #4223, CST), Bax (1:1000, #5023, CST), and β-actin (1:1000, # Ab5694, Abcam). After three 10-min TBST washes, membranes were incubated with species-matched HRP-conjugated secondary antibodies (anti-rabbit: 1:2,000, # Ab7074, Abcam; anti-mouse: 1:2,000, # Ab7076, Abcam) for 1 h at 25°C. Membranes underwent three 5-min TBST washes before chemiluminescent detection using ECL Prime substrate (WBKLS0500; Millipore) with 5-min substrate incubation. β-actin served as loading control. Protein signals were digitally captured using a ChemiDoc XRS+ system (12003154, Bio-Rad).

### 2.9 Co-immunoprecipitation analysis

Collect cells and add 1mL IP lysis buffer. After full lysis, centrifuge at 4 degrees and 12,000g for 15 minutes to obtain total cell protein as the input group. Use 20µl magnetic beads (#P2151, BeyoMag, Inc.) to immunoprecipitate the lysate and incubate with anti-LAMP2A antibody (1:1000, #Ab125068, Abcam) at 4°C overnight. Samples were centrifuged (250×g, 5 min, 4°C), and supernatants removed via vacuum aspiration. Magnetic beads were washed thrice with ice-cold PBS. Following three PBS washes, immunocomplex-bound beads were resuspended in 60 μL IP lysis buffer combined with 15 μL 2× Laemmli sample buffer, then thermally denatured at 100°C for 10 min using a programmable heating block (ThermoFisher, 88870001). The supernatant is transferred to a new centrifuge tube, and the resulting product is the IP product, which is stored at -20 degrees. Western blot assay was performed using an anti-DJ-1 antibody (dilution 1:1000, # Ab76008, Abcam).

### 2.10 Statistical methods

Statistical analyses were performed using SPSS Statistics 23.0 (v23.0.0; IBM). For two-group comparisons, independent two-sample Student’s t-tests with Welch’s correction were applied. Multigroup analyses employed one-way ANOVA followed by Dunnett’s post hoc test for comparisons against a specified control group. *p<0.05 indicates statistical significance.

## 3. Results

### 3.1 Upregulation of LAMP2A in Gastric Cancer Promotes CMA

First, we validated the expression of LAMP2A in GC using immunohistochemistry in gastric tumors and normal tissue (Fig. 1A). Then, we analyzed the transcript levels of LAMP2A in paired gastric adenocarcinoma and adjacent normal mucosa from the TCGA and GEO databases (Fig. 1B). Four gastric adenocarcinoma cell lines and non-neoplastic gastric epithelial cells were analyzed. LAMP2A mRNA expression was quantified by RT-PCR (Fig. 1C). Protein levels were assessed via immunoblotting WB (Fig. 1D). Our analysis revealed that LAMP2A upregulation in gastric adenocarcinoma tissues versus matched normal mucosa. both transcript and protein levels of LAMP2A were increased in gastric cancer cells. LAMP2A is the rate-limiting component of chaperone-mediated autophagy (CMA), which directly governs lysosomal substrate flux through its assembly into multimeric translocation complexes. Up-regulated and down-regulated expression of LAMP2A can enhance and decrease the activity of CMA, respectively [15]. Therefore, the activity of CMA can be determined by detecting the level of LAMP2A. Compared with normal gastric mucosa cells, CMA activity was enhanced in gastric cancer cells.

**Fig. 1.**
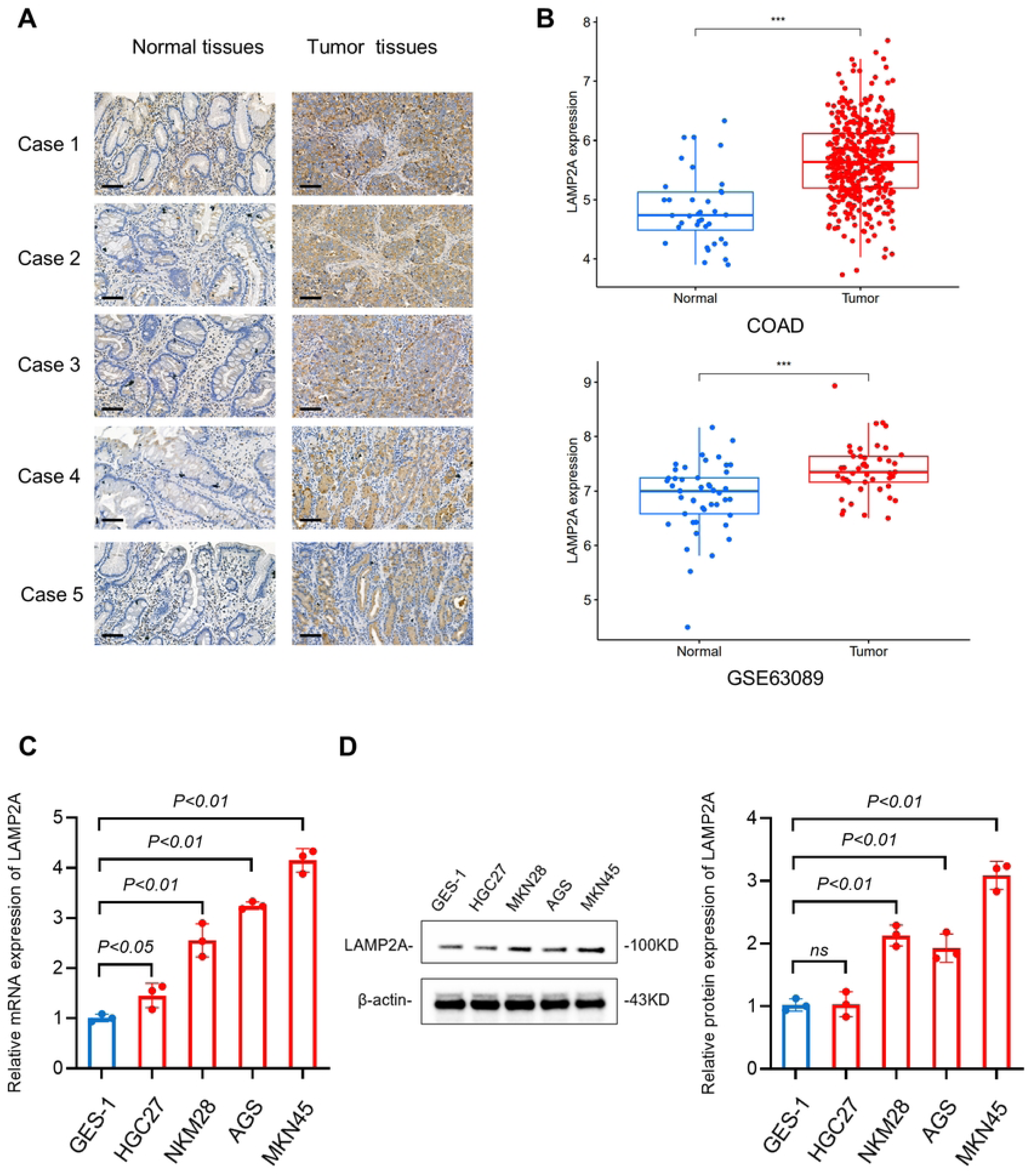
Detection of LAMP2A expression in gastric cancer. (A) Immunohistochemical staining of LAMP2A in gastric tumor and normal tissue. (Scale bar: 20 μm). (B) The TCGA and GEO databases show the mRNA expression levels of LAMP2A in gastric tumors and normal tissue. (C)The qPCR results showed the LAMP2A expression at mRNA levels. (D)Western blot analysis of the level of LAMP2A in several cell lines. Note:TCGA(https://portal.gdc.cancer.gov/)GEO(https://www.ncbi.nlm.nih.gov/geo/query/acc.cgi? acc=GSE63089). Data are presented as the mean ± SD. n = 3, ns, no significance, *P<0.05, **P<0.01 and ***P<0.001.

### 3.2 Oxidative stress up-regulated the expression of LAMP2A in gastric cancer cells

Experimental evidence indicates that oxidative stress induces LAMP2A upregulation in glioblastoma cells [16]. Hydrogen peroxide (H_2_O_2_) treatment has been established as a standard method for generating experimental models of oxidative stress [17]. To investigate oxidative stress as an inducer of LAMP2A upregulation in gastric cancer, four cell lines were exposed to H_2_O_2_ at two concentrations of 150µM and 300µM, respectively, to establish a cell model of severe oxidative stress. Western blot analysis demonstrated H₂O₂-induced elevation of LAMP2A protein levels. It was suggested that oxidative stress was a driver of LAMP2A upregulation in GC (Fig. 2).

**Fig. 2.**
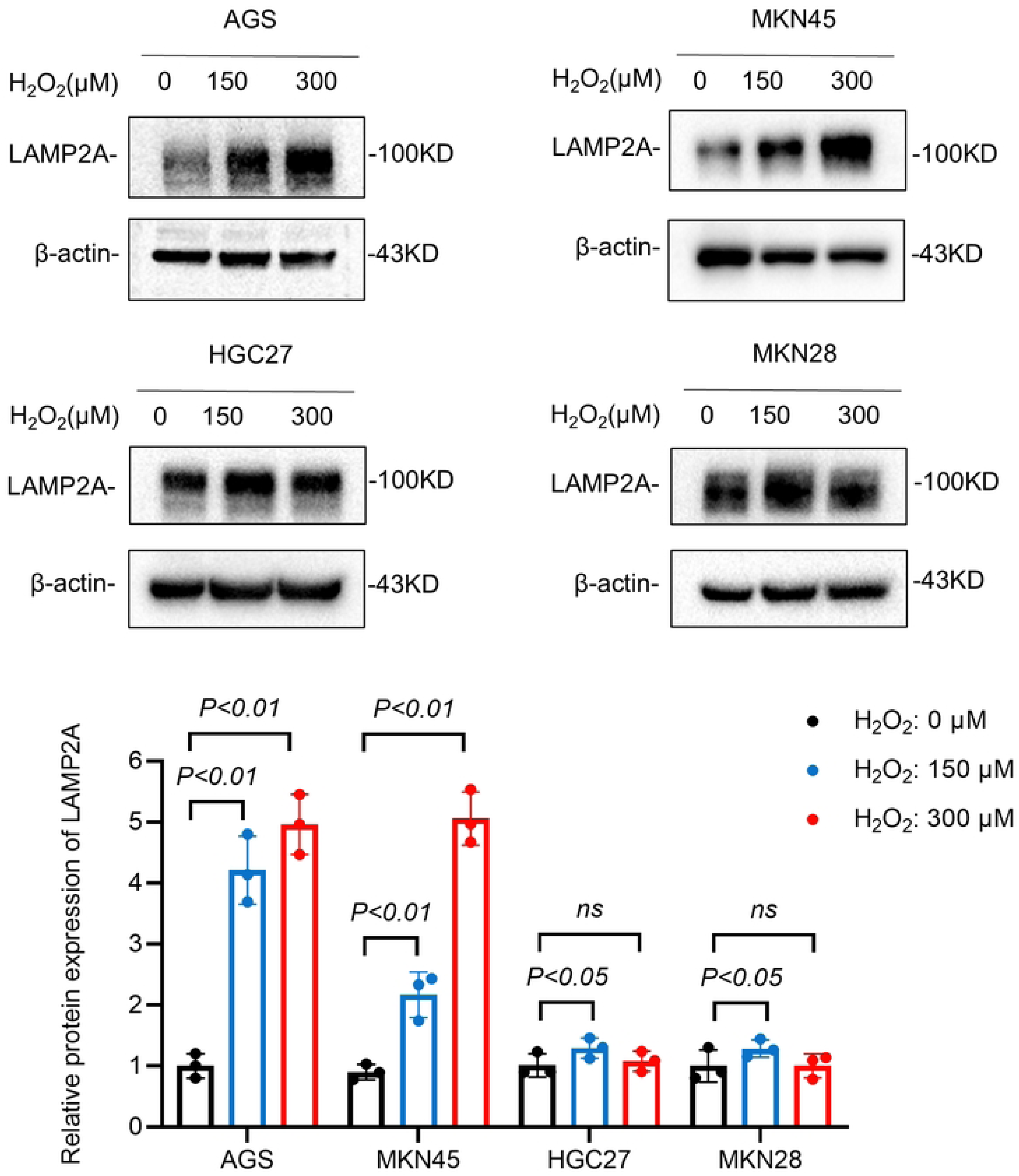
Oxidative stress upregulates the expression of LAMP2A in cell lines. Four cell lines were exposed to H_2_O_2_ (150µM, 300µM), respectively, to establish a severe oxidative stress cell model. WB results showed the protein expression of LAMP2A.

### 3.3 Construction of LAMP2A knockdown and overexpression cell models

AGS cells were transfected with a recombinant plasmid encoding LAMP2A to establish stable overexpression, with experimental groups designated as: negative control (NC) and LAMP2A-overexpressing (LAMP2A). MKN45 cells transduced with LAMP2A-targeted shRNA lentiviral particles to create knockdown models, with groups designated as: negative control (LV-sh NC) and LAMP2A-knockdown (LV-sh L2A).

RT-qPCR showed a significant decrease of LAMP2A in MKN45 cells compared to the LV-sh NC control (p<0.01, Fig. 3A). LAMP2A mRNA expression was significantly elevated in AGS cells compared with negative control (NC) cells (**p < 0.01; Fig. 3C). Western blot analysis confirmed successful LAMP2A knockdown and overexpression (*p < 0.05, **p < 0.01; Fig. 3B, D). The LAMP2A downregulation and overexpression models were successfully constructed and can be used for subsequent functional experiments.

**Fig. 3.**
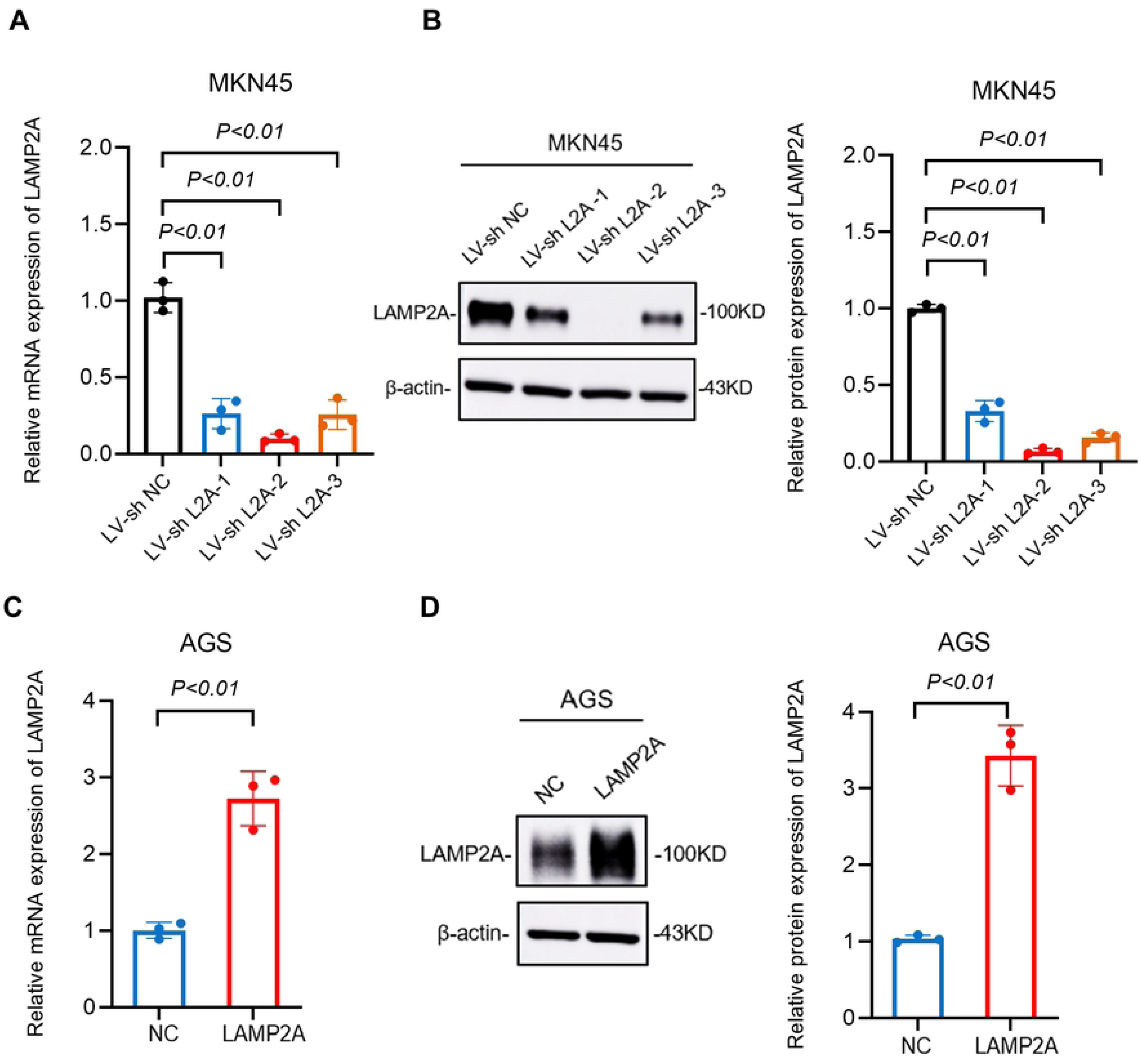
Validation of LAMP2A knockdown and overexpression efficiency in cellular models through qPCR and WB. (A, B) Verification of LAMP2A knockdown efficiency in MKN45 cell. (C, D) Verification of LAMP2A overexpression efficiency in AGS cell. Note: n = 3, *p < 0.05, **p < 0.01. NC, negative control.

### 3.4 Loss of LAMP2A promoted oxidative stress-induced apoptosis of gastric cancer cells

To investigate the functional role of LAMP2A in oxidative stress response, we used the previously constructed LAMP2A cell model. CCK-8 analysis revealed that LAMP2A knockdown significantly impaired cellular proliferative capacity compared to controls (Fig. 4A). Downregulation of LAMP2A also inhibited proliferation when cells were stimulated with 150µM H_2_O_2_. We also demonstrated a significant amplification of proliferation rate disparity between LAMP2A and control groups under H_2_O_2_-induced oxidative stress relative to basal culture conditions (Fig. 4B). Flow cytometric analysis revealed that LAMP2A-knockdown gastric cancer cells exhibited dose-responsive increases in early apoptosis with 150 μM and 300 μM H2O2 exposure (Fig. 4C). Conversely, the early apoptosis rate of the LAMP2A-overexpressing cells was significantly lower than that in the control group (Fig. 4D). These findings demonstrate that LAMP2A knockdown inhibit CMA activity, thereby sensitizing gastric cancer cells to oxidative stress-induced apoptosis.

**Fig. 4.**
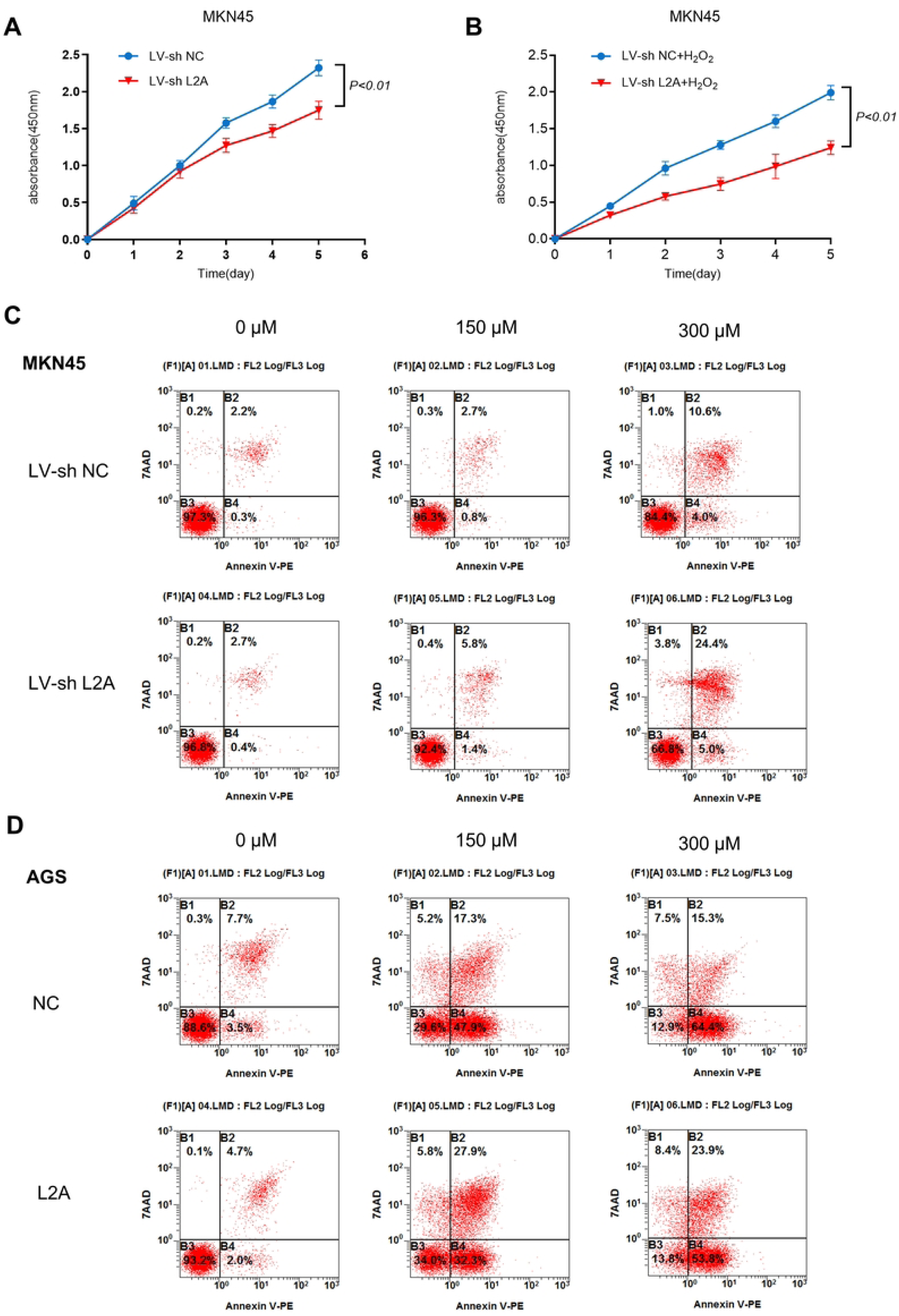
LAMP2A knockdown promotes apoptosis of gastric cancer cells induced by severe oxidative stress. (A): Proliferation of the control group and LAMP2A-knockdown group. (B): Proliferation of the control group and LAMP2A-knockdown group following 150 μM H_2_O_2_ treatment. (C): LAMP2A knockdown cell lines were exposed to H_2_O_2_ (0, 150µM and 300µM), and the apoptosis rate was measured by FACS after 24-hour treatment. D: LAMP2A over-expressing cells were exposed to H_2_O_2_ (0, 150µM and 300µM), and the apoptosis rate was measured by FACS after 24-hour treatment. The data represent the mean ± standard deviation from three independent experiments. *P<0.05. L2A, LAMP2A; NC, negative control.

### 3.5 LAMP2A Recognizes DJ-1 as a Chaperone-Mediated Autophagy Substrate Through Direct Interaction

Upregulation of chaperone-mediated autophagy (CMA) rate-limiting component LAMP2A in gastric cancer correlates with CMA activation during tumor progression. Experimental evidence demonstrates that CMA sustains tumor cell proliferation via targeted degradation of RND3. The antioxidant protein DJ-1 (Parkinson disease protein 7, PARK7) may be a substrate for CMA [13]. DJ-1 (PARK7) suppresses the apoptosis signal-regulating kinase 1 (ASK1) pathway under oxidative stress, attenuating stress-induced apoptosis and protecting cells from oxidative damage [18]. CMA substrate selectivity requires recognition of a conserved pentapeptide motif (KFERQ-like sequence) in target proteins [19]. Structural analysis reveals that DJ-1 contains an evolutionarily conserved KFERQ-like motif (Fig. 5A), establishing it as a CMA substrate through this recognition signature. WB revealed elevated DJ-1 protein levels in gastric cancer cell lines, with further upregulation observed under hydrogen peroxide treatment (Fig. 5B, C). To investigate DJ-1’s role in LAMP2A-mediated (CMA) under oxidative stress conditions, we assessed LAMP2A-DJ-1 interactions via co-immunoprecipitation (Co-IP) and immunofluorescence (IF), comparing basal and H₂O₂-treated conditions. Co-IP results demonstrated enhanced DJ-1–LAMP2A binding following 24-hour H₂O₂(300μM) treatment compared to untreated controls (Fig. 5D). Immunofluorescence imaging confirmed marked colocalization of LAMP2A and DJ-1 under oxidative stress versus basal conditions (Fig. 5E).

**Fig. 5.**
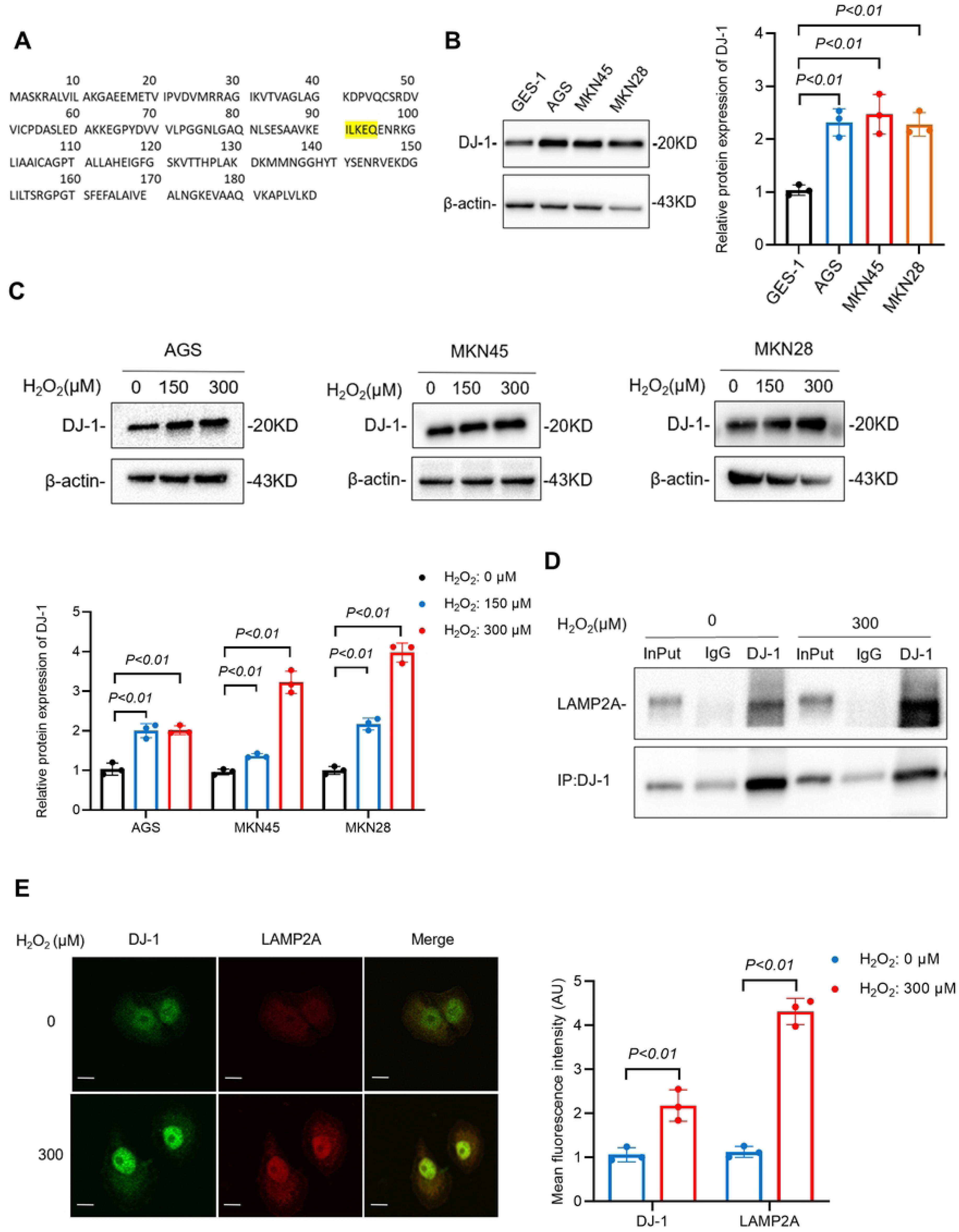
DJ-1 is the substrate for CMA, and LAMP2A directly interacts with DJ-1. (A): Identification of a conserved KFERQ-like motif in DJ-1, a CMA substrate recognition signature. (B): WB analysis of DJ-1 protein expression levels. (C) H₂O₂ dose-dependent upregulation of DJ-1 protein in gastric cancer cells (150 μM, 300 μM; 24 h treatment). (D): Co-IP confirms enhanced DJ-1/LAMP2A interaction in H₂O₂-treated (300 μM, 24 h) MKN45 cancer cells. (E): IF demonstrates oxidative stress-induced colocalization of DJ-1 and LAMP2A in MKN45 cells (300 μM H₂O₂, 24 h).

### 3.6 LAMP2A knockdown promotes DJ-1-induced apoptosis of gastric cancer cells under severe oxidative stress

To explore whether LAMP2A could regulate the expression of DJ-1, we constructed the LAMP2A knockdown gastric cancer cell line MKN45shL2A and its control cell line MKN45shNC, treated gastric cancer cells with hydrogen peroxide (0, 300µM) for 24 hours, and detected the changes of DJ-1 after LAMP2A knockdown by WB. The results showed that LAMP2A knockdown increased DJ-1 expression regardless of H_2_O_2_ stimulation. WB analysis demonstrated that LAMP2A knockdown-mediated DJ-1 upregulation significantly elevated pro-apoptotic BAX expression while suppressing anti-apoptotic Bcl-2 levels, indicative of a pro-apoptotic shift in cellular signaling (Fig. 6).

**Fig. 6.**
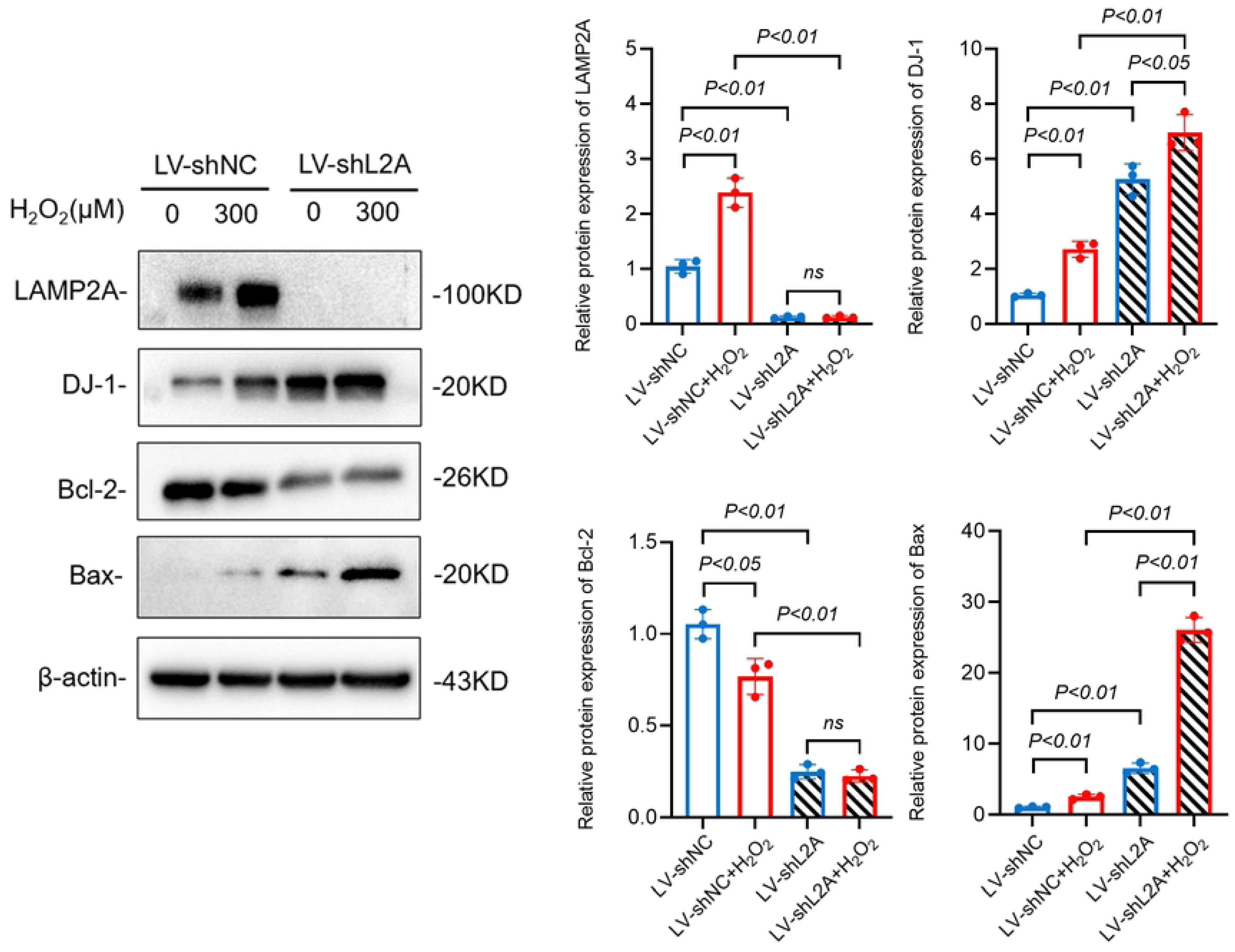
LAMP2A knockdown promotes DJ-1-induced apoptosis of gastric cancer cells under severe oxidative stress. MKN45shL2A and MKN45shNC were stimulated by H_2_O_2_ (0,300 µm) for 24 hours, and the protein levels of LAMP2A, DJ-1, apoptosis-related proteins Bcl-2 and BAX were detected by WB.

**Fig. 7.**
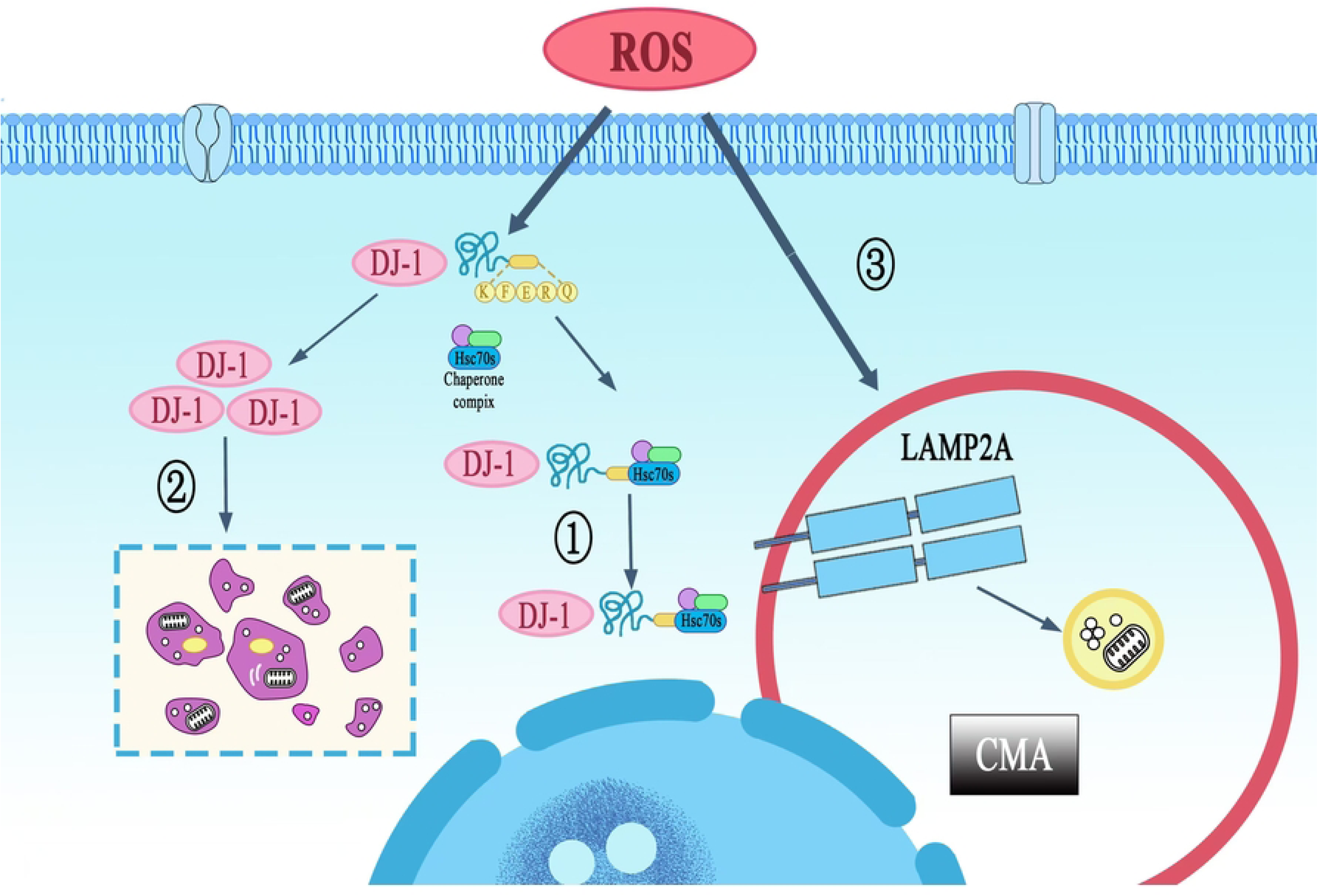
Mechanisms of LAMP2A-dependent CMA enhances oxidative stress resistance. ①CMA inhibits tumor cell apoptosis through selective removal of overoxidized DJ-1 protein. ②In pathological conditions, ROS-mediated overoxidation converts DJ-1 into a pro-apoptotic form that triggers gastric cancer cell death, distinct from its physiological oxidative modification under basal oxidative stress. ③Oxidative stress serves as the primary inducer of LAMP2A upregulation in gastric cancer cells, establishing a compensatory mechanism for CMA activation.

### 3.7 Mechanisms of LAMP2A-dependent CMA enhances oxidative stress resistance in gastric cancer cells

## 4. Discussion

Autophagy, a conserved lysosomal degradation pathway, preserves cellular homeostasis by selectively removing damaged organelles and misfolded proteins. It comprises three subtypes: macroautophagy (cytoplasmic cargo engulfed by double-membrane autophagosomes for lysosomal fusion), microautophagy, and chaperone-mediated autophagy (CMA). In contrast, microautophagy involves direct lysosomal membrane invagination, where substrates are internalized via vesicle formation and engulfed for proteolytic processing [20]. CMA is mechanistically distinct from macroautophagy and microautophagy. Substrate proteins are selectively recognized by cytosolic chaperones, directly translocated across the lysosomal membrane via LAMP2A receptors, and degraded without intermediate vesicle encapsulation.

CMA is the first autophagy discovered to be selective for protein degradation. It regulates cell function by specifically recognizing an amino acid sequence “KFERQ” in substrate proteins, thereby selectively degrading key intracellular proteins [21]. Emerging evidence indicates that CMA is activated across multiple tumor types. Tumor microenvironmental stress conditions, including nutrient deprivation, hypoxia, and elevated reactive oxygen species (ROS), serving as potent inducers of CMA activity [22]. Therefore, investigating the relationship between CMA and oxidative stress resistance in tumor cells, and elucidating the underlying molecular mechanisms, holds significant research value.

LAMP2A is a key receptor molecule in the CMA process. Substrate proteins require binding to LAMP2A on the lysosomal membrane for translocation into the lysosomal lumen and subsequent degradation. Increased LAMP2A expression enhances chaperone-mediated autophagy (CMA) activity, with its protein abundance serving as a direct quantitative indicator of CMA functionality [23]. Previous studies demonstrated that the expression of LAMP2A was significantly elevated in gastric cancer [14], indicating that the basic activity of CMA in tumors was enhanced. It has been reported that ROS can upregulate the level of LAMP2A in glioblastoma [10]. Building on the observed LAMP2A upregulation in gastric malignancies, we hypothesized its potential role in mediating oxidative stress adaptation. To test this, four established gastric adenocarcinoma cell lines (AGS, MKN45, HGC27, MKN28) were exposed to H_2_O_2_ gradients for 24h. WB analysis demonstrated dose-dependent elevation of LAMP2A protein abundance under oxidative stress conditions (Fig. 2). These findings establish oxidative stress as a potent inducer of LAMP2A expression in gastric cancer cells. To delineate the mechanistic role of LAMP2A in oxidative stress adaptation, we establish oxidative stress models, gastric adenocarcinoma cells were exposed to H₂O₂ gradients. LAMP2A-knockdown and LAMP2A-overexpression cell lines were generated. Cellular responses were quantified using CCK-8 proliferation assays and FACS. The results showed that loss of LAMP2A inhibited CMA activity and increased the sensitivity of gastric cancer cells to oxidative stress.

Studies have reported that the oxidative stress regulatory protein DJ-1 (PARK7) may bind to LAMP2A and HSC70, suggesting that they are substrates of CMA [14]. DJ-1 is a new antioxidant molecule discovered in the past decade. During oxidative stress, it can regulate the redox signaling pathway in cells and induce the production of other antioxidant molecules [24]. Tumor cells can also use these antioxidant mechanisms to protect themselves from damage by ROS. DJ-1 is elevated in many malignant tumors and promotes tumor progression. For example, in primary breast cancer and primary lung cancer, DJ-1 inhibits the function of the tumor suppressor PTEN, increases the phosphorylation level of PTEN’s downstream target PKB/AKT, and enhances the activity of PKB/AKT, thereby promoting tumor survival [25]. Elevated DJ-1 expression in colorectal malignancies drives tumorigenic progression through enhanced proliferative, migratory and invasive capacities, with clinico-pathological analyses confirming its correlation with advanced TNM staging and reduced 5-year overall survival [26,27]. In pancreatic cancer, papillary thyroid cancer, and osteosarcoma, DJ-1 knockout inhibits tumor proliferation and promotes apoptosis [28,29]. Emerging evidence in gastric carcinogenesis demonstrates DJ-1 promotes gastric cancer resistance, participates in metastasis, and is associated with poor prognosis [30,31]. These collective findings establish DJ-1 as a critical cytoprotective molecule of tumor cell.

We initially demonstrated that DJ-1 protein expression levels were elevated in gastric cancer cells following oxidative stress induction, showing its cytoprotective function through redox regulation. However, Co-IP assays revealed enhanced DJ-1/LAMP2A complex formation under oxidative stress relative to basal conditions. IF analysis further demonstrated pronounced subcellular co-localization of these proteins during severe oxidative stress (Fig 5). These findings indicate selective degradation of a DJ-1 subpopulation via CMA. Given DJ-1’s established cytoprotective properties through redox homeostasis maintenance and apoptosis inhibition. There is an apparent paradox-CMA targets DJ-1 for degradation despite its canonical cytoprotective role.

Accordingly, we have undertaken research to address this paradox. Some studies have found that in tumor cells, physiological oxidation enables DJ-1 to sequester ASK1 through direct protein-protein interaction, thereby suppressing JNK/p38 signaling cascades while potentiating cytoprotective autophagy flux. Conversely, pathological hyperoxidation induces DJ-1-ASK1 complex dissociation, liberating ASK1 to phosphorylate downstream effectors including p38 MAPK that ultimately execute apoptosis commitment [32]. This indicates that hyperoxidized DJ-1 is cytotoxic, and cells should clear hyperoxidized DJ-1 in time to facilitate normal survival. Meanwhile, Studies have found that the proteasome inhibitor MG132 cannot significantly increase the content of DJ-1 [33], suggesting that its degradation depends on the lysosomal system.

Therefore, we propose a cytoprotective paradigm wherein oxidative stress-induced DJ-1/LAMP2A complex formation facilitates CMA-mediated selective degradation of hyperoxidized DJ-1 isoforms, thereby enabling apoptosis evasion through maintenance of redox-competent DJ-1 pools in malignant cells. Therefore, we verified this hypothesis by using the constructed LAMP2A knockdown cell model, and the results showed that down-regulation of LAMP2A could increase the protein level of DJ-1. In addition, we also found that LAMP2A depletion precipitated hyperoxidized DJ-1 accumulation concomitant with apoptotic priming, evidenced by reciprocal BAX upregulation and Bcl-2 downregulation (Fig 6). These results indicate that hyperoxidized DJ-1 is degraded through the CMA pathway, which provides a basis for the conclusion that hyperoxidized DJ-1 is the substrate protein of CMA.

Normally, hyperoxidation of DJ-1 occurs exclusively during severe oxidative stress induced by H_2_O_2_ exposure. LAMP2A knockdown under these conditions compromises CMA-mediated clearance of hyperoxidized DJ-1, resulting in increase of DJ-1. Notably, in the absence of H_2_O_2_ stimulation, we also found that LAMP2A knockdown induced DJ-1 accumulation. Regarding this phenomenon, we believe that under physiological conditions, CMA operates as a housekeeping mechanism that sustains proteostasis through continuous clearance of endogenously generated hyperoxidized DJ-1, thereby maintaining redox equilibrium under physiological conditions. Current evidence indicates that CMA may regulate neuronal mitochondrial homeostasis by degrading oxidatively damaged DJ-1 protein, as demonstrated in a recent study [34]. However, whether DJ-1 undergoes CMA-dependent degradation remains controversial, with insufficient experimental evidence to conclusively resolve this question. Our findings in gastric cancer models provide experimental evidence supporting the classification of overoxidized DJ-1 as a CMA substrate in tumors. A key methodological constraint in this study involves our inability to directly interrogate the oxidation state of DJ-1. While we inferred its overoxidized status through hydrogen peroxide-induced oxidative stress, this approach precludes definitive attribution of apoptotic effects specifically to DJ-1 hyperoxidation, lacking evidence that over-oxidized DJ-1 could directly induce apoptosis. Thus, the mechanistic role of CMA in counteracting oxidative stress during gastric carcinogenesis requires systematic investigation. Furthermore, more studies are needed to definitively establish DJ-1 hyperoxidation as a biomarker of CMA substrate specificity in malignant contexts.

## 5. Conclusions

Our study found that elevated expression of LAMP2A and DJ-1 in gastric cancer cell lines compared to normal gastric mucosal cells, with oxidative stress identified as an inducible factor driving their upregulation. Mechanistically, we found hyperoxidized DJ-1 as a chaperone-mediated autophagy (CMA) substrate, demonstrating that CMA-mediated clearance of oxidized DJ-1 enhances cellular antioxidant capacity and promotes gastric cancer cell survival. These findings elucidate the mechanistic role of chaperone-mediated autophagy (CMA) in regulating tumor cell redox homeostasis. This work provides a molecular framework for advancing diagnostic stratification and molecularly targeted therapeutic strategies in gastric cancer.

## List of abbreviations

ATCC: American Type Culture Collection
BCA: Bicinchoninic acid
CCK-8: Cell Counting K it-8
GC: Gastric cancer
DAPI: 4’,6-Diamidino-2-phenylindole
DMEM: Dulbecco’s modified Eagle’s medium FBS Fetal bovine serum
H_2_O_2_: Hydrogen peroxide
NC: Negative control
PBS: Phosphate buffered solution
qPCR: Quantitative polymerase chain reaction
shRNA: Short hairpin RNA
WB: Western blot

## Acknowledgements

Not applicable.

## Availability of data and materials

The datasets generated and/or analyzed during the current study are not publicly available due to privacy of laboratory content but are available from the corresponding author on reasonable request.

## Authors’ contributions

Conceptualization, MGP, SSL; methodology, TTG, SSL; investigation, MGP, GTT and SSL; analysis and interpretation of data, YZ, TTG and TJT; writing–original draft, MGP, SSL; writing—review and editing, MGP; acquisition of funding and supervision of the research, ZY and MGP. All authors have read and agreed to the published version of the manuscript.

## Ethics approval and consent to participate

Not applicable.

## Patient consent for publication

Not applicable.

## References

1. Bray, Freddie, Ferlay, Jacques, Soerjomataram, Isabelle, Siegel, Rebecca L., Torre, Lindsey A. and Jemal, Ahmedin. Global Cancer Statistics 2018: GLOBOCAN Estimates of Incidence and Mortality Worldwide for 36 Cancers in 185 Countries. CA: A Cancer Journal for Clinicians, 2018. 68(6): 394–424.

2. Wen-Long Guan, Ye He, and Rui-Hua Xu. Gastric cancer treatment: recent progress and future perspectives. Journal of Hematology & Oncology. 2023. 16(57).1–28.

3. Yuqing Ren, Ruizhi Wang, Siyuan Weng, Hui Xu, Yuyuan Zhang, Shuang Chen, Shutong Liu, Yuhao Ba, Zhaokai Zhou, Peng Luo, Quan Cheng, Qin Dang, Zaoqu Liu1 and Xinwei Han. Multifaceted role of redox pattern in the tumor immune microenvironment regarding autophagy and apoptosis. Molecular Cancer. 2023. 22(130).1–24.

4. Yujie Geng, Zhuo Wang, Jiaying Zhou, Mingguang Zhu, Jiang Liu and Tony D. James. Recent progress in the development of fluorescent probes for imaging pathological oxidative stress. Chem. Soc. Rev. 2023. 52. 3873–3926.

5. Yuqi Wang, Jingli Xu, Zhenjie Fu, Ruolan Zhang Weiwei Zhu, Qianyu Zhao, Ping Wang, Can Hu, Xiangdong Cheng. The role of reactive oxygen species in gastric cancer. Cancer Biol Med 2024. 0182 1–14.

6. Jayanta Debnath, Noor Gammoh and Kevin M. Ryan. Autophagy and autophagy-related pathways in cancer. Nature Reviews Molecular Cell Biology 24, 560–575 (2023).

7. João Vasco Ferreira, Ana da Rosa Soares, José Ramalho, Catarina Máximo Carvalho, Maria Helena Cardoso, Petra Pintado, Ana Sofia Carvalho, Hans Christian Beck, Rune Matthiesen, Mónica Zuzarte, Henrique Girão, Guillaume van Niel and Paulo Pereira. LAMP2A regulates the loading of proteins into exosomes. SCIENCE ADVANCES 8, eabm1140 (2022).

8. Qi Jia, Jin Li, Xiaofeng Guo, Yi Li, You Wu, Yuliang Peng, Zongping Fang, Xijing Zhang. Neuroprotective effects of chaperone-mediated autophagy in neurodegenerative diseases. NEURAL REGENERATION RESEARCH 19(6):1291–1298 (2024).

9. Ruchen Yao, Jun Shen. Chaperone-mediated autophagy: Molecular mechanisms, biological functions, and diseases. MedComm 4(5), e347 (2023).

10. Alessia Lo Dico, Daniela Salvatore, Cristina Martelli, Dario Ronchi, Cecilia Diceglie, Giovanni Lucignani and Luisa Ottobrini. Intracellular Redox-Balance Involvement in Temozolomide Resistance-Related Molecular Mechanisms in Glioblastoma. Cells 8(11):1315 (2019).

11. Ying Xuan, Shuang Zhao, Xingjun Xiao, Liwei Xiang and Hua-Chuan Zheng. Inhibition of chaperone-mediated autophagy reduces tumor growth and metastasis and promotes drug sensitivity in colorectal cancer. MOLECULAR MEDICINE REPORTS 23: 360, (2021).

12. Rui-Fang Dong, Cheng-Jiao Qin, Yong Yin, Liang-Liang Han, Cheng-Mei Xiao, Kai-Di Wang, Rong-Yuan Wei, Yuan-Zheng Xia and Ling-Yi Kong. Discovery of a potent inhibitor of chaperone-mediated autophagy that targets the HSC70–LAMP2A interaction in non-small cell lung cancer cells. British Pharmacological Society bph.16165 (2023).

13. Tapas Saha. LAMP2A overexpression in breast tumors promotes cancer cell survival via chaperone-mediated autophagy. Autophagy 8(11), 1643–1656 (2012).

14. Jinfeng Zhou, Jianjun Yang, Xing Fan, Sijun Hu, Fenlizhou, Jiaqiang Dong, Song Zhang, Yulong Shang, Xiaoming Jiang, Hao Guo, Ning Chen, Xiao Xiao, Jianqiu Sheng, Kaichun Wu, Yongzhan Nie and Daiming Fan. Chaperone-mediated autophagy regulates proliferation by targeting RND3 in gastric cancer. Autophagy 12(3):515–528(2016).

15. Jaione Auzmendi-Iriarte and Ander Matheu. Intrinsic role of chaperone-mediated autophagy in cancer stem cell maintenance. Autophagy 18(12):3035–3036 (2022).

16. Helmut Sies and Dean P. Jones. Reactive oxygen species (ROS) as pleiotropic physiological signalling agents. Nature Reviews Molecular Cell Biology 21(7):363–383 (2020).

17. Alessia Lo Dico, Daniela Salvatore, Cristina Martelli, Dario Ronchi, Cecilia Diceglie, Giovanni Lucignani and Luisa Ottobrini. Intracellular Redox-Balance Involvement in Temozolomide Resistance-Related Molecular Mechanisms in Glioblastoma. 8(11):1315 (2019).

18. Stephanie E. Oh and M. Maral Mouradian. Cytoprotective mechanisms of DJ-1 against oxidative stress through modulating ERK1/2 and ASK1 signal transduction. Redox Biology 14:211–217 (2018).

19. Susmita Kaushik and Ana Maria Cuervo. The coming of age of chaperone-mediated autophagy. Nature Reviews Molecular Cell Biology 19(6):365–381 (2018).

20. Fernanda V. Duraes, Jennifer Niven, Juan Dubrot, Stéphanie Hugues and Monique Gannagé. Macroautophagy in endogenous processing of self- and pathogen derived antigens for MHC class ii presentation. Frontiers in Immunology 6:459 (2015).

21. Mathieu Bourdenx, Evripidis Gavathiotis and Ana Maria Cuervo. Chaperone-mediated autophagy: a gatekeeper of neuronal proteostasis. Autophagy 17(8):2040–2042 (2021).

22. Shuangshuang Le, Xin Fu, Maogui Pang, Yao Zhou, Guoqing Yin, Jie Zhang and Daiming Fan. The Antioxidative Role of Chaperone-Mediated Autophagy as a Downstream Regulator of Oxidative Stress in Human Diseases.21:1–15 (2022).

23. Lei Qiao, Jiayi Hu, Xiaohan Qiu, Chunlin Wang, Jieqiong Peng, Cheng Zhang, Meng Zhang, Huixia Lu, and Wenqiang Chen. LAMP2A, LAMP2B and LAMP2C: similar structures, divergent roles. Autophagy 19(11):2837–2852 (2023).

24. Line Duborg Skou, Steffi Krudt Johansen, Justyna Okarmus and Morten Meyer. Pathogenesis of DJ-1/PARK7-Mediated Parkinson’s Disease. Cells 13(4):296 (2024).

25. Raymond H. Kim, Malte Peters, YingJu Jang, Wei Shi, Melania Pintilie, Graham C. Fletcher, Carmela DeLuca, Jennifer Liepa, Lily Zhou, Bryan Snow, Richard C. Binari, Armen S. Manoukian, Mark R. Bray, Fei-Fei Liu, Ming-Sound Tsao and Tak W. Mak. DJ-1, a novel regulator of the tumor suppressor PTEN. Cancer Cell 7(3):263–73(2005).

26. Xiaojian Zhu, Chen Luo, Kang Lin, Fanqin Bu, Fan Ye, Chao Huang, Hongliang Luo, Jun Huang, Zhengming Zhu. Overexpression of DJ-1 enhances colorectal cancer cell proliferation through the cyclin-D1/MDM2-p53 signaling pathway. BioScience Trends 14(2):83–95 (2020).

27. Jing Zhou, Hao Liu, Lian Zhang, Xin Liu, Chundong Zhang, Yitao Wang, Qing He, Ying Zhang, Yi Li, Quanmei Chen, Lu Zhang, Kui Wang, Youquan Bu and Yunlong Lei. DJ-1 promotes colorectal cancer progression through activating PLAGL2/ Wnt/BMP4 axis. Cell Death and Disease 9(9):865 (2018).

28. Kai Qiu, Qingji Xie, Shan Jiang and Ting Lin. Silencing of DJ-1 reduces proliferation, invasion, and migration of papillary thyroid cancer cells in vitro, probably by increase of PTEN expression. International Journal of Clinical and Experimental Pathology 12(6):2046–2055 (2019).

29. Zhanjun Ma, Jingjing Yang, Yang Yang, Xuexi Wang, Guohu Chen, Ancheng Shi, Yubao Lu, Shouning Jia, Xuewen Kang, Li Lu. Rosmarinic acid exerts an anticancer effect on osteosarcoma cells by inhibiting DJ-1 via regulation of the PTEN-PI3K-Akt signaling pathway. Phytomedicine 68:153186 (2020).

30. Lejia Qiu, Zhaoxia Ma, Xiaoran Li, Yizhang Deng, Guangling Duan, Le Zhao, Xingwang Xu, Lin Xiao, Haoyue Liu, Zhengming Zhu, and Heping Chen. DJ-1 is involved in the multidrug resistance of SGC7901 gastric cancer cells through PTEN/PI3K/Akt/Nrf2 pathway. Acta Biochim Biophys Sin 52(11):1202–1214 (2020).

31. Xue-kai PAN, Fei SU, Li-hua XU, Zhang-shuo YANG, Dan-wen WANG, Li-jie YANG, Fanzheng KONG, Wei XIE, Mao-hui FENG. DJ-1 Alters Epirubicin-induced Apoptosis via Modulating Epirubicin-activated Autophagy in Human Gastric Cancer Cells. Current Medical Science 38(6):1018–1024 (2018).

32. Ji Cao, Meidan Ying, Nan Xie, Guanyu Lin, Rong Dong, Jun Zhang, Hailin Yan, Xiaochun Yang, Qiaojun He and Bo Yang. The Oxidation States of DJ-1 Dictate the Cell Fate in Response to Oxidative Stress Triggered by 4-HPR: Autophagy or Apoptosis? ANTIOXIDANTS & REDOX SIGNALING 21(10):1443–1459 (2014).

33. Karin Gorner, Eve Holtorf, Sabine Odoy, Brigitte Nuscher, Ayako Yamamoto, Jorg T. Regula, Klaus Beyer, Christian Haass, and Philipp J. Kahle. Differential Effects of Parkinson’s Disease-associated Mutations on Stability and Folding of DJ-1. THE JOURNAL OF BIOLOGICAL CHEMISTRY 279(8):6943–6951 (2004).

34. Bao Wang, Zhibiao Cai, Kai Tao, Weijun Zeng, Fangfang Lu, Ruixin Yang, Dayun Feng, Guodong Gao and Qian Yang. Essential Control of Mitochondrial Morphology and Function by Chaperone-mediated Autophagy through Degradation of PARK7. Autophagy 12(8):1215–1228 (2016).

